# Enabling population protein dynamics through Bayesian modeling

**DOI:** 10.1101/2023.10.30.564713

**Authors:** Sylvain Lehmann, Jérôme Vialaret, Audrey Gabelle, Luc Bauchet, Jean-Philippe Villemin, Christophe Hirtz, Jacques Colinge

**Author notes:** Correspondance: Prof. Jacques Colinge.

## Abstract

The knowledge of protein dynamics or turnover in patients provides invaluable information related to certain diseases, drug efficacy, or biological processes. A great corpus of experimental and computational methods has been developed, including by us, in the case of human patients followed *in vivo*. Moving one step further, we propose here a new modeling approach to capture the highly relevant notion of population protein dynamics. Using two data sets, we show that models inspired by population pharmacokinetics can accurately capture protein turnover within a cohort of individuals, even in presence of substantial inter-individual variability. Such models pave the way for comparative studies searching for altered dynamics or biomarkers in diseases.

## INTRODUCTION

Beyond the knowledge of protein abundance in various tissues (Meyer and Schilling, 2017), there is great interest in learning about the dynamics of proteins (Doherty and Whitfield, 2011). Protein dynamics, which is commonly referred to as protein turnover, is the net rate at which proteins are produced or imported in a tissue, and simultaneously degraded or cleared. This complementary and dynamic perspective to protein abundance is relevant in a number of applications of clinical proteomics. For example, in different pathologies, abnormal turnover has been observed for specific proteins such as amyloid-β (A*β*), Tau, or sTREM2 in Alzheimer disease (AD) (Mawuenyega *et al*., 2010; Sato *et al*., 2018; Suárez-Calvet *et al*., 2016), retinol-binding protein 4 (RBP4) in diabetes (Jourdan *et al*., 2009), or tissue remodeling during early-stage human heart failure (Lam *et al*., 2014). Besides clinical applications, protein turnover can be related to fundamental biological processes such as heart morphogenesis (Konzer *et al*., 2013). Turnover data are typically acquired by mass spectrometry (MS) after introducing isotopic tracers to label the newly synthesized proteins (Bateman *et al*., 2006; Jaleel *et al*., 2006; Doherty *et al*., 2012; Claydon *et al*., 2012; Wilkinson, 2018). The ratio of labeled versus unlabeled protein peptide abundances is called the relative isotope abundance (RIA). The variation of RIA over time provides turnover information.

Different protocols can be followed regarding the introduction of a tracer, *e*.*g*., through the diet, intravenous injections, or even the medium if we consider cells in culture or organoids. Our interest lies in human – or animal – *in vivo* studies, where biofluids represent the most convenient and ethically acceptable material for sequential measures on a same individual. Following an initial publication of our labeling protocol (Lehmann *et al*., 2015), we developed a flexible and accurate 2-compartment mathematical model (Lehmann *et al*., 2019). In particular, we showed that this general model was able to fit data obtained by stable isotope labeling kinetics (SILK) (Bateman *et al*., 2006). SILK is a pulse-chase protocol, where ^13^C_6_-Leu is intravenously injected for nine hours, allowing to observe new protein synthesis, but also clearance, by acquiring sequential samples over many hours, 24 hours for instance. The new work we present here is a follow-up bringing the modeling, and the extraction of kinetic parameters, to the population level. Namely, given a cohort or population of individuals that were submitted to SILK, we want to learn the typical values, variability, and correlation of the protein dynamics parameters over the whole population. This type of mathematical model is common in population pharmacokinetics (Bauer *et al*., 2007), where compound availability and clearance in patients must be characterized population-wide to adjust standard regimens. In the case of protein turnover, our perspective is to provide robust and comprehensive models of healthy homeostatic states, in comparison to disease states. This knowledge could obviously lead to progresses in biomarker discovery as well as diagnostic applications beyond pure research and protein classification.

Similar to population pharmacokinetics studies, we developed hierarchical a Bayesian model whose parameters were fit with Markov-chain Monte-Carlo (MCMC) sampling. The new model is illustrated on an unpublished cohort of seven individuals whose blood plasma samples were analyzed with targeted MS, *i*.*e*., multiple reaction monitoring (MRM). To contrast this first cohort that displayed moderate inter-individual variability, we exploited a second cohort of four individual cerebrospinal fluid (CSF) samples. The later were not completely comparable and thus provided much more heterogeneous inter-individual data, which enabled us to challenge our population model robustness.

## MATERIALS AND METHODS

### Human samples

Samples were generated following the clinical protocol “In Vivo Alzheimer Proteomics (PROMARA)” (ClinicalTrials Identifier: NCT02263235), which was authorized by the French ethical committee CPP Sud-Méditerranée IV (#2011-003926-28) and by the ANSM agency (#121457A-11). Enrolled patients (group a) were hospitalized in neurosurgery unit due to subarachnoid hemorrhage and received a temporary ventricular derivation of the CSF. The experiment protein turnover started 8 to 19 days after initial, medical ventricular drainage and normalization of CSF clinical chemistry analysis (normal CSF protein content lies in the 0.2-0.4 g/L range (Roche *et al*., 2008)). Additional patients (group b) were hospitalized in neurology in relation with cognitive impairment etiologic investigation. Patient data are reported in Table S1. CSF and blood plasma were collected at multiple time points after injection of the tracer for roughly 24 (CSF) or up to 36 (plasma) hours. We applied the ethically approved (see above) original SILK ^13^C_6_-Leu infusion protocol (Bateman *et al*., 2006). Briefly, ^13^C_6_-Leu prepared *per* the European Pharmacopeia [19] was intravenously administered. After a 10 min initial bolus at 2 mg/kg, an 8h50 infusion at 2 mg kg/h was performed. Ventricular CSF or plasma EDTA samples were collected starting at the beginning of the ^13^C_6_-leucine infusion, roughly every 3h (3 to 6 mL). Samples were transported to the laboratory at 4°C, and centrifuged at 2000g for 10 minutes. CSF and plasma samples were aliquoted into 1.5-mL polypropylene tubes and stored at –80°C until further analysis.

In this study, we analysed CSF samples from four patients of group a (Pat1a to Pat4a) and seven of group b (Pat7b-Pat13b). Patients were selected based on availability of CSF and plasma MS samples at multiple time points.

### Sample analysis

Sample preparation was automated on AssayMap BRAVO (Agilent T., Santa Clara, US) to reduce preanalytical variability. Briefly, 2μL of plasma or 30uL of CSF were used. Protein samples were reduced and alkylated, and digested with trypsin prior to LC-MS analysis.

The MRM protocol was reported in previous publications (Percy *et al*., 2013; Hirtz *et al*., 2018; Lehmann *et al*., 2019); we hence only summarize the main steps here. Proteins were selected for their relevance to neurodegenerative diseases and clear detection in previous, proteome-wide experiments in plasma and CSF by our laboratory. The reporter peptides were selected for their high signal intensity in these previous experiments. Tables S2 and S3 reports the selected proteins and peptides for CSF and plasma samples. MRM was executed on the samples directly using a 1290 liquid chromatography (LC) system (Agilent Technologies) equipped with a reverse-phase column (RRHD Eclipse Plus C18) coupled with a QqQ MS instrument (6490, Agilent technologies). The MS instrument worked in dynamic MRM with a retention time window of 4.5 minutes and a maximum cycle time fixed at 700ms. All the analyses were performed in duplicates. A minimum of one peptide *per* protein and 3 transitions by peptide were required. Skyline 4.1 was used to process raw MS data. Figure 1A presents an overview of the LC and MS pipeline.

**Figure 1.**
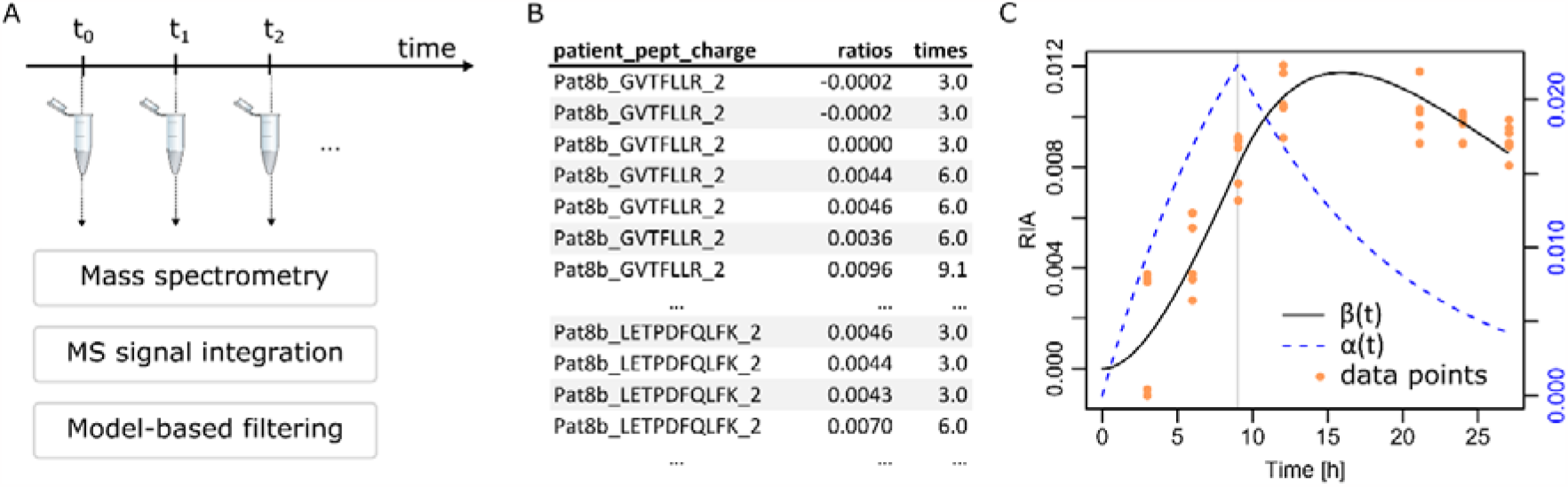
Simplified workflow. (A) Samples were collected at multiple time points and analyzed by MS. Integration of all the spectra found for a given peptide at a given time point was performed with Skyline. Filtering of peptides for which sufficient and interpretable signals were available was performed employing a mathematical model of protein turnover. (B) Representative data for a given protein (A1BG), for which two different peptides were followed in MRM and both measured with double charges. Corresponding acquisition times and RIA (ratios) were available. (C) A1BG ratios (orange dots) along with the fitted mathematical model. The vertical gray line indicates the end of tracer injection at 9 hours.

Although they were not the object of this work, we also used two samples (CSF and plasma from PatA1) that were subjected to the proteome-wide (non MRM) protocol as describe in our previous publication (Lehmann *et al*., 2019). The CSF sample was discussed in this latter publication, whereas the blood sample remains unpublished. The data generated were subjected to the data analysis workflow described below. We obtained reliable turnover data for roughly 200 proteins in each sample, whose parameters were only used to learn the typical range of parameter values for mathematical modeling (see below) from diverse proteins.

### MS data processing and existing individual mathematical model

Integrated MS spectra by Skyline were the input for fitting protein dynamics models. Typical input data are illustrated in Figure 1B. The processing of these data has been described in details already (Lehmann *et al*., 2019), we hence provide a summary only. First, we briefly reintroduce our turnover model. For a given peptide and a given time, the observed RIA (Figure 1B) is defined by the ratio of the heavy Leu signal *P*_*H*_ (observed at a shifted mass of +6 Da *per* Leu) and the total signal *P*_*L*_ + *P*_*H*_, *P*_*L*_ the signal at the nominal mass. We denote the curve of RIA over time by *β*(*t*). Our 2-compartment model relies on a notion of rate of tracer availability (first compartment) denoted *α*(*t*) to be involved in protein synthesis (second compartment). To parallel pharmacokinetic 2-compartment models, *α*(*t*) relates to the ratio of initial drug concentration divided by the volume of distribution. Note that in the case of protein turnover, only ratios are modeled and hence *α* and *β* are dimensionless. They are related by the following ordinary differential equations (ODEs):

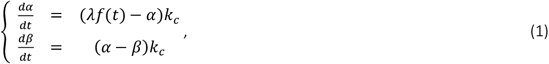

with *α*(0) = 0 = *β*(0). Figure 1C illustrates a typical dataset with the *α* and *β* curves. It is important to observe that λ, which relates to tracer availability for protein synthesis, essentially acts as a scale parameter, whereas *k*_*c*_, the clearance/degradation rate primarily acts as a shape parameter that conditions protein half-life. Due to the generally large ratio of intensities between *P*_*L*_ and *P*_*H*_, we have shown in that noise causes an almost uniform vertical shift of the observed RIA values (Lehmann *et al*., 2019). Therefore, we proposed an algorithm that include the computation of an optimal shift along with the parameters λ and *k*_*c*_ to adjust *β*(*t*) to the data. Summed squared errors with respect to observed RIAs were weighted proportionally to 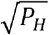 (RIAs with stronger *P*_*H*_ signals were more accurate). Weighted summed squared errors were minimized by a quasi-Newton iteration (function optim in R with method BFGS) to adjust the parameters. Eq. (1) was numerically integrated by a stiff Runge-Kutta method (Hairer and Wanner, 1996).

To achieve robust results in the presence of noisy RIAs, parameter fitting was iterative with a first application of the above to call outlier RIAs. RIAs were considered outliers provided they lied at a distance larger than half the difference between the minimum and maximum values of the first fitted *β*(*t*) model. A second application of the quasi-Newton method without the outliers was then performed. In addition, RIAs at time 0 were always considered outliers since no tracer incorporation had occurred yet. Our original data processing pipeline finished with the application of a bootstrap to estimate confidence intervals. Here, we used the R library boot to perform a nonparametric balanced bootstrap (100 times), while the original publication used a parametric Gaussian bootstrap. In the sequel, we refer to the parameter and CI95 estimates found by this procedure as QNB for quasi-Newton-bootstrap.

## RESULTS AND DISCUSSION

### Initial Bayesian models

We started the construction of a Bayesian population model by first establishing a Bayesian model equivalent to our QNB original procedure presented above since it led to accurate models. Minimization of the differences between observed RIAs and model-predicted values is achieved by the computation, and maximization of a likelihood in the Bayesian paradigm. Assuming RIA_*i*_ to be the *i*^th^ observation and *β*_*i*_ the corresponding model value, we naturally have

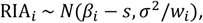

with *i* ∈ {1; ⋯; *n*} and *n* the number of RIAs, *β*_*i*_ = *β*(*t*_*i*_), *t*_*i*_ the time at which RIA_*i*_ was observed (due to replicates, several *t*_*i*_ can be identical with different indices *i*), *w*_*i*_ the weight proportional to 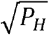 for observation *i*, and *s* the vertical shift to acknowledge the noise in the ratios (Materials and Methods). *N*(*a, b*^*2*^) denotes a normal distribution with mean *a* and variance *b*^*2*^.

To obtain a complete, hierarchical Bayesian model, we introduce prior distributions on the model parameters as well as the mean and variance of the error normal distribution. The resulting model is the following:

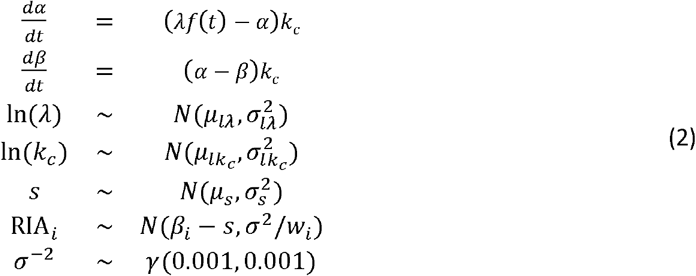

We denote by γ(*a, b*) a Gamma distribution with shape *a* and rate *b*. The prior parameters for *s*, In(*k*_*c*_), and In(λ) were learned from a large number of models (roughly 200 distinct proteins) fitted with the QNB algorithm using unbiased data, *i*.*e*., not MRM data (see Materials and Methods, and Tables S2 & S3). In the case of *s*, we set μ_*s*_ = 0, and 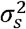 at 10^3^ for the plasma samples and 1/500 for the slightly more noisy CSF data. The Gamma prior for σ^−*2*^, *i*.*e*., the precision, is a commonly used vague (non-informative) prior.

We implemented the MCMC sampling for the parameters *σ, s*, In(*k*_*c*_), and In(λ) in the above model using two approaches, both implemented in R. First, we defined a function proportional to the log probability density of (2) and used the R libraries mcmc and adaptMCMC. Alternatively, we used BUGS to define the model and OpenBUGS (Lunn *et al*., 2000) through its R interface R2OpenBUGS. Some particular *β*(*t*) shapes such as apolipoprotein A1 (APOA1) in CSF illustrated in Fig2A were more difficult to fit, and adaptMCMC and mcmc failed to find correct parameters (Figure S1). OpenBUGS managed to address those more difficult data efficiently. In the majority cases, typically illustrated by neuropilin-2 (NRP2) and complex component 1s (C1S) in Figure 2A, the three library produced almost identical parameter estimations (Figure S1). The distribution-aware OpenBUGS Gibbs sampler was thus more effective generally, and we decided to use OpenBUGS only. The BUGS code and the R function for adaptMCMC and mcmc are provided in SI. We found that 100 000 iterations including 50 000 burn-in were sufficient for OpenBUGS safe convergence. We systematically used two Markov chains, and a convergence diagnostic was achieved comparing within- and between-chain variability (Brooks and Gelman, 1998). Each chain was initialized with random λ, and *k*_*c*_ values drawn from their respective prior distributions. The shift *s* was initialized at 0 and σ^*2*^ at 0.1.

**Figure 2.**
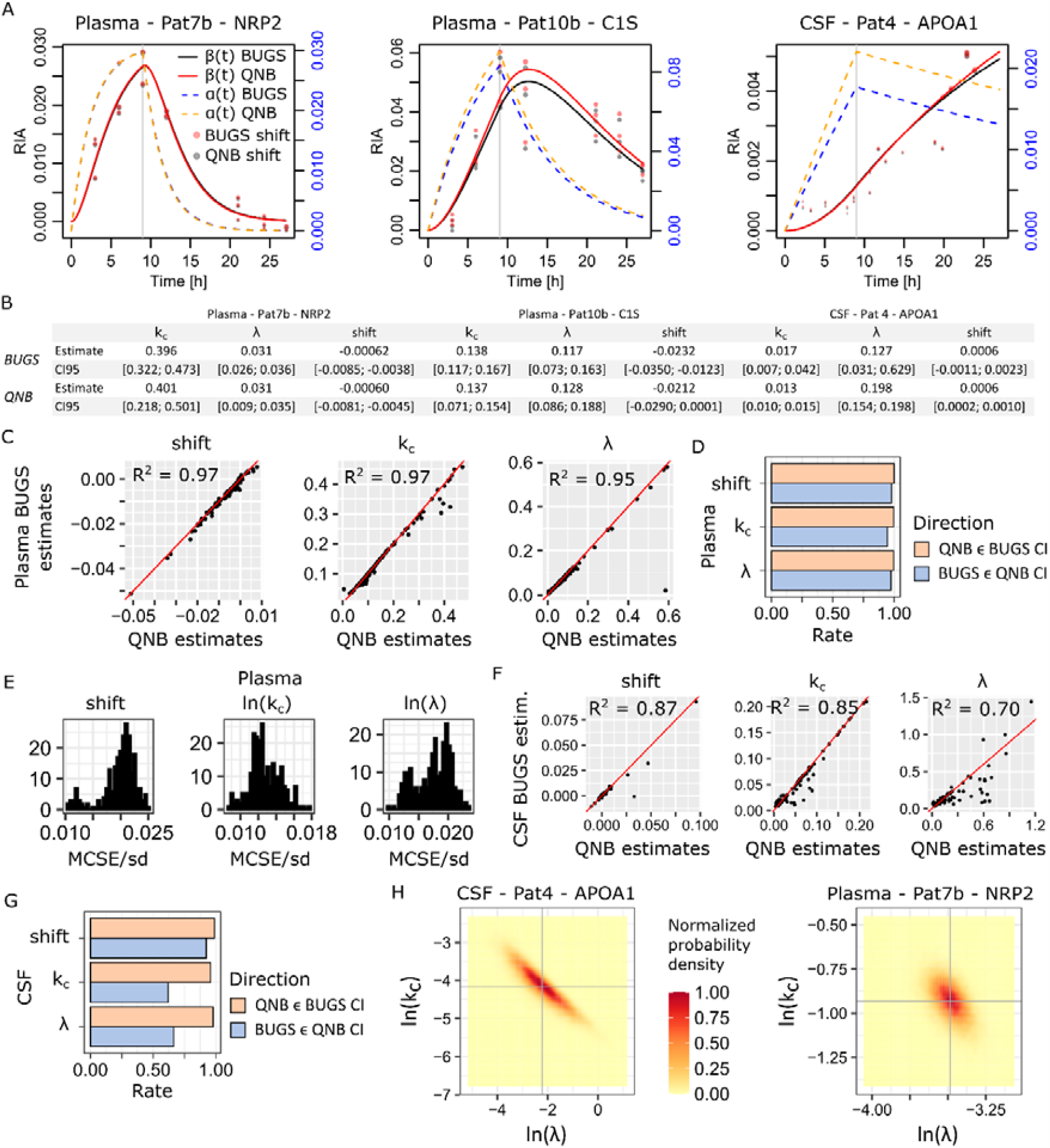
Initial Bayesian models of single proteins in single patients. (A) Three representative examples comparing the BUGS fitted models with the original QNB solutions. (B) Comparison of the BUGS *versus* QNB fitted parameters shift, *k_c_* and λ. Spearman squared correlation is denoted *R* ^2^. (C) Global comparison of BUGS versus QNB estimated parameters over all the plasma proteins and patients. (D) Rate of inclusion of the QNB parameter estimates in the BUGS 95% credible intervals, and conversely. (E) Autocorrelation-corrected MCSE/estimate standard deviation distributions. (F) BUGS *versus* QNB parameter estimate correlation over all the CSF proteins and patients. (G) Inclusion rates of QNB estimates in BUGS 95% credible intervals and *vice versa*. (H) MCMC sampled probability density over the (λ, *k_c_*)-space. Marginal means are featured by the gray crosses.

Outliers identified by the QNB algorithm were removed from the data given to OpenBUGS (same for mcmc and adaptMCMC) since the latter was sensitive to some extreme outliers (data not shown). In theory, it is possible to replace the normal distribution for errors on RIA_*i*_ in Eq. (2) by a heavy-tailed distribution such as Student with degree of freedom 4 to address outliers. Nonetheless, here our goal was not to replace the satisfying QNB algorithm, but to build a population model on the top of it. We thus trusted QNB outlier calling and exploited it.

From Figure 2A, we see that the Bayesian solution is usually close to QNB, NRP2 and C1S being representative of the majority of the cases. The inferred parameters for the two algorithms are reported in Figure 2B with estimated 95% confidence intervals (QNB) and 95% credible intervals (BUGS). Indeed, comparing the parameter values for all the plasma proteins in patients Pat7b to Pat13b (37 different proteins, 236 individual proteins in total, Table S3), we found an excellent correlation between the original QNB estimates and those from OpenBUGS (Figure 2C). We also found high compatibility between the 95% confidence and credible intervals (Figure 2D). As a commonly accepted rule of thumb, the number of iterations in a MCMC estimation should be such that the MCMC standard error (MCSE) divided by the standard deviation of the sampled parameter should be below 5%. As reported in Figure 2E, this was achieved for the three estimated parameters using a conservative value for MCSE that was corrected for autocorrelation (OpenBUGS “Time-series SE” estimates). This validated the choice of the number of iterations. Considering CSF data (Pat1 to Pat4, 26 different proteins, 92 individual proteins in total, Table S4), we made similar observations (Figure 2F), though with lower correlation. This is explained by the fact that in CSF data, *β*(*t*) shapes similar to APOA1 in Figure 2A represented roughly half the data. Credible intervals estimated by OpenBUGS were reliable and almost always included QNB estimates, while the converse was rather in the 70% range, but for the shift that was highly compatible with QNB estimations. MCSE/sd values were similar to Figure 2E.

To understand better the CSF results, we first observed that in the typical example of APOA1 (Figure 2A), the curve *β*(*t*) was properly fit to the experimental RIAs by both algorithms. Nevertheless, the QNB and BUGS *α*(*t*) curves were different. This indicated a stronger correlation between the model parameters λ and *k*_*c*_ that manifested by the ability to compensate variation in one by the value of the other. Indeed, comparing the probability density estimated by MCMC sampling over the (λ, *k*_*e*_)-space, we found a large, almost linear area for CSF Pat4 APOA1 and a narrower, rounder area for plasma Pat7b NRP2 (Figure 2H). The latter configuration is typical of easy and fast convergence, whereas the former indicates a more difficult and slower convergence. This was also reflected in BUGS credible intervals sizes in Figure 2B. The Bayesian approach did a better job at estimating realistic 95% credible intervals, almost always including QNB estimates. This is not the case for the 95% confidence intervals estimated by QNB, which tended to be too narrow including no more than ∼70% of the BUGS estimates. Lastly, it is important to remember that in our model in Eq. (1), the most relevant, shape- or half-life-related parameter is *k*_*c*_ for which a rather high *R*^*2*^ = 0.85 (Spearman) was achieved.

### Population hierarchical Bayesian model

To develop a population model, we start with plasma data, typical examples are featured in Figure 3A (the 27 plasma proteins for which we had data for every patient are depicted in Figure S2).

**Figure 3.**
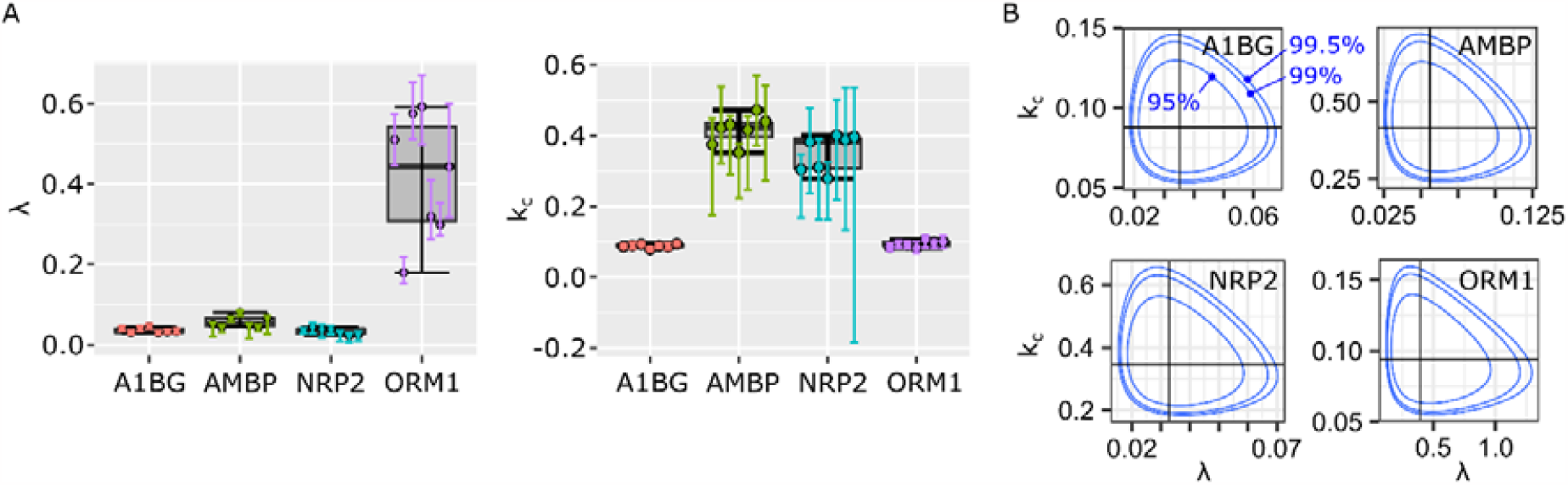
Four example proteins in plasma data. (A) Variability of individual parameters for the seven patients. Dots represent QNB estimates, the vertical color bars their respective CI95s, and a boxplot computed form the individual estimates was added to the background to suggest population dispersion. (B) Areas covered in the (λ, *k_c_*) parameter space by the true Bayesian population models. Concentric boundaries indicate the space occupied by 95% (inside the innermost boundary), 99% and 99.5% of the population as estimated from the available cohort.

The principle of population Bayesian modeling consists in adding a population level to the model in Eq. (2). This additional layer should capture parameter typical values as well as potential correlations between them (Figure 2H). If we define the parameter vector

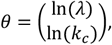

then this is achieved by a 2-dimensional normal distribution

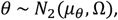

with mean μ_θ_ and variance Ω. Hyper-priors are introduced for these two quantities

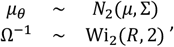

where the vector *μ* ∈ ℝ^*2*^ components are respectively set to the means of In(λ) and In(*k*_*e*_). Those means were obtained from a large set of previous observations as we did for the individual models above (Tables S3-4). For Σ, we employed a commonly used vague hyper-prior with

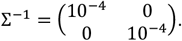

Wi^*2*^(*R*, ^*2*^) denotes a 2-dimensional Wishart distribution that generalizes the Gamma distribution to multidimensional variates (compare Ω^−1^ above with σ^−*2*^ in Eq. (2)). A largely used vague hyper-prior is obtained with

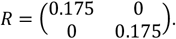

Lastly, writing RIA_*ki*_ the *i*^th^ observed RIA for patient *k*, the residuals likelihood is defined by

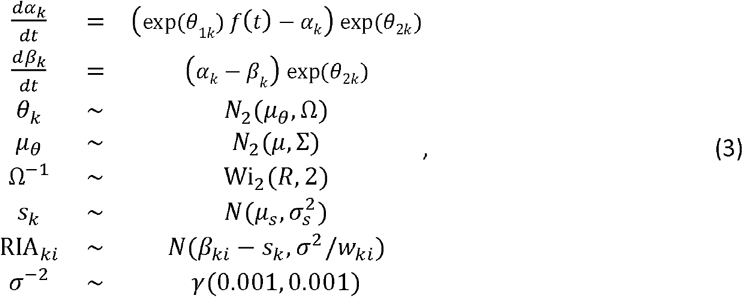

with (similar to the individual model) *β*_*ki*_ the value of *β*_*k*_(*t*) at the time where RIA_*ki*_ was observed, and the corresponding weight *w*_*ki*_. Then, θ_1*k*_ is In(λ_*k*_) and θ^*2*^_*k*_ is In(*k*_*ck*_), the logarithms of patient *k* turnover parameters.

As we did for the individual models, we ran the BUGS model (reported in SI) for 100 000 iterations including 50 000 burn-ins, using two chains. The Markov chains were initialized with random *θ*_*k*_ drawn from their prior, *s*_*k*_ = 0, and *σ*^*2*^ = 0.01. Convergence was reached for all the proteins but alpha-1-microglobulin/bikunin precursor (AMBP), beta-2-microglobulin (B2M), C1S, and NRP2. Running the BUGS models, same data and initial chains, with WinBUGS that offers real-time visual tracking, we found that *θ* components were diverging in opposite directions. One parameter became indefinitely large, and the other one indefinitely small to compensate. We could easily solve these four cases by either starting the two chains with the parameter values θ_*k*_ set to the QNB estimates or, alternatively, by using a prior *N*^*2*^(*μ*, Σ) with *μ* and Σ learned from the seven QNB estimates available for each protein specifically (*μ* was set to the average, and diagonal elements of Σ^−1^ to 1/variances). Applying these two alternative procedures to the 24 proteins for which the initial approach worked, we found no real differences in the learned population models (Figure S3).Moreover, AMBP, B2M, C1S, and NRP2 shared the same fast turnover dynamics, which was apparently difficult for the initial approach. Based on the comparison of the three approaches (Figure S2), we concluded that data were strong enough to make the choice of a specific prior consequence less. We hence obtained *bona fide* population parameters for all the considered 27 plasma proteins. Figure 3B illustrates the population models for the example proteins in Figure 3A. In particular, Figure 3B shows the population probability density over the parameter space (λ, *k*_*c*_). We note that the additional parameter space area covered switching from 95% to 99 % or 99.5% of the population is very small indicating a robust estimate of turnover diversity for each protein.

Another benefit of a population model is to provide us with the ability to estimate the range of protein dynamics that can be expected by 95% of the population, or any percentage of interest. This is simply achieved generating a large sample of turnover parameter values from the population model *N*^*2*^(*μ*_*θ*_, Ω). In Figure 4, four examples of protein dynamic range are featured along with the individual curves fit from the same population model (Eq. (3)), but using the *θ*_*k*_’s. Figure S4 features AMBP, B2M, C1S, and NRP2 dynamics similarly to show again the absence of real impact of the prior choice.

**Figure 4.**
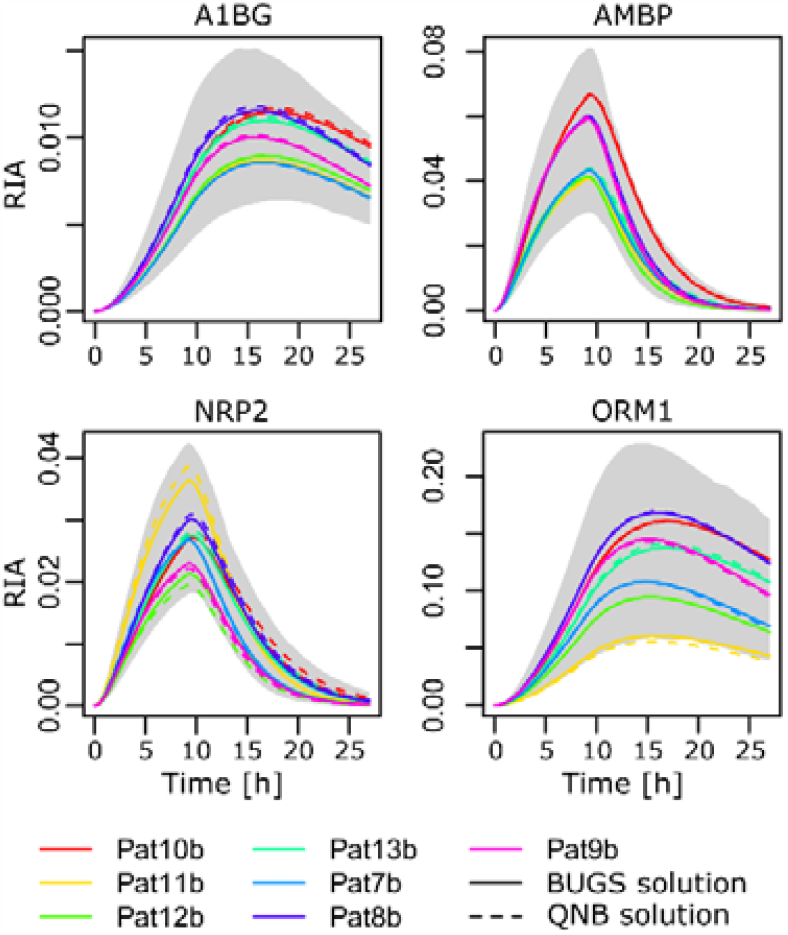
Population dynamics in plasma samples. Areas in the (time, RIA)-space that are covered by 95% of the population as estimated by our model based on the available cohort. The gray 95% area was obtained by generating 500 (λ, *k_c_*) pairs and computing the CI95 of all the 500 resulting *β*(*t*) curves at each time point. The individual curves for the BUGS model were obtained from the population model (Eq. (3)) using patient-specific *θ*_*k*_’s. They were as accurate as the solutions found by individual BUGS model using Eq. (2) and illustrated in Figure 2A.

### Population modeling of heterogeneous data

CSF protein data displayed a much higher heterogeneity between patients compared to the plasma data set. This variability was due to that fact that the four CSF patients experienced subarachnoid hemorrhage, which introduced blood in the CSF. Part of their treatment included a drainage of ventricular CSF, which provided the opportunity to collect the samples. Although these samples were collected at a late stage of the therapy, assessing that protein concentration in CSF returned to a normal range (Table S1), we cannot consider the CSF patients as really comparable. In this methodological work, where no biological conclusion is made, this offers us the opportunity to confront our Bayesian approach with much more variable data.

Out of the 19 proteins for which we had data for all four patients (from group B), our initial approach (random initial chains and *θ* prior parameter trained on a set of roughly 200 diverse proteins, Table S3) could fit 12/19 proteins. Compared to 4/27 difficult case in plasma, we note the expected additional difficulty. Next, we tried the two alternative procedures applied to the difficult cases in plasma, *i*.*e*., initial chains on the QNB estimates or more informative prior. Only the informative prior approach resulted in an improvement, which we interpret as an indication that the problem was really difficult. Even accurate starting points led to divergence. Only a restriction of the search space through the prior helped. We could obtain 18/19 models, and AMBP (in CSF) remained impossible to model. Figure 5A illustrates different CSF cases covering all configurations of heterogeneity on signal intensity (λ heterogeneity) and turnover (*k*_*c*_ heterogeneity), see Figure S5 for all the 18 models. Figure 5B shows population coverage of the parameter space. Figure 5C explains why CSF AMBP was so reluctant to population modeling, and Figure 5D illustrates population dynamics. We note that our population model was successful in pretty heterogeneous cases such as A2M, and to a certain extent TTR, in Figure 5D.

**Figure 5.**
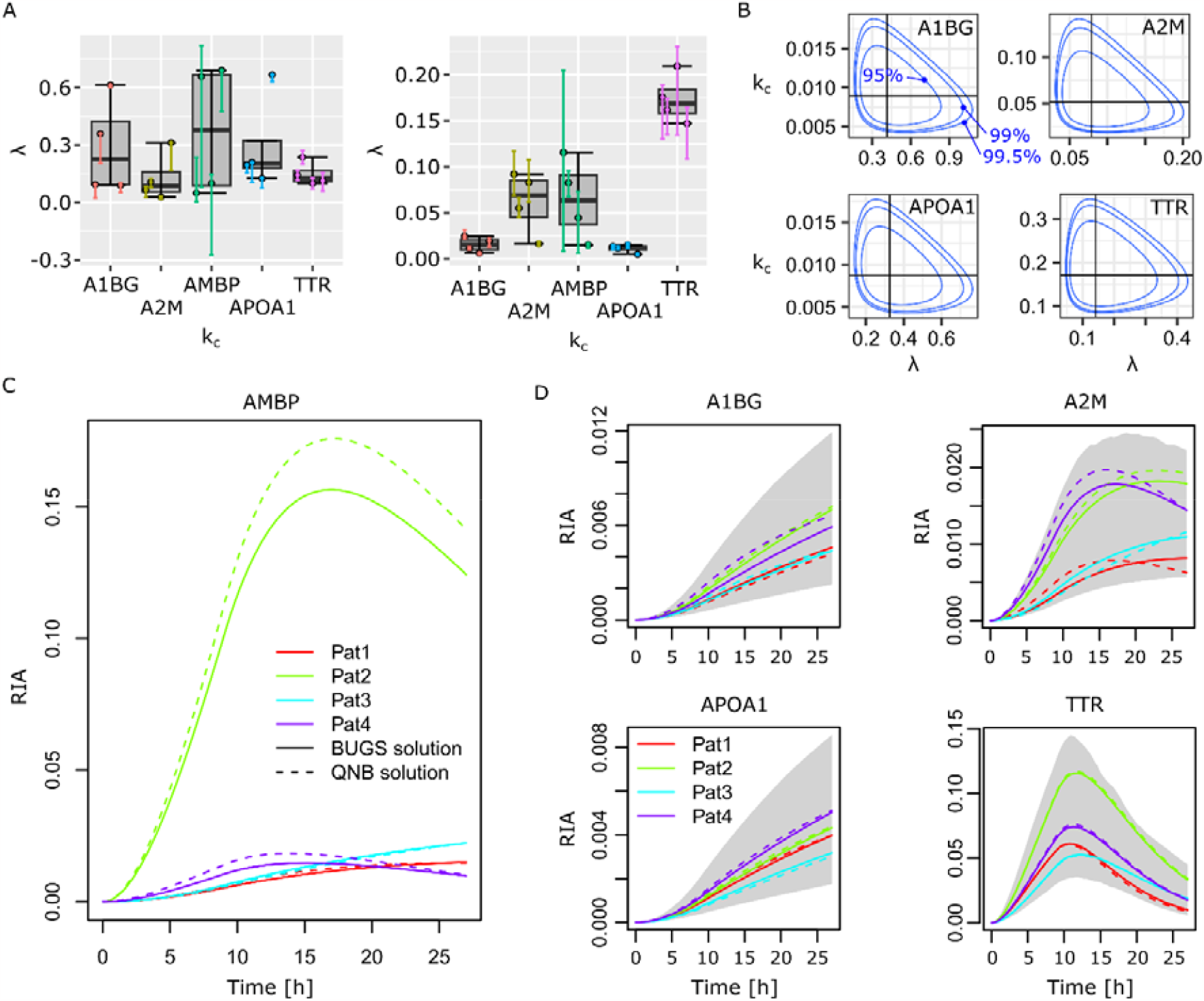
Modeling heterogeneous dynamics. (A) Individual parameter variability in the four CSF patients from group a. (B) Areas in the Areas covered in the (λ, *k_c_*) parameter space by the true Bayesian population models. Concentric boundaries indicate the space occupied by 95% (inside the innermost boundary), 99% and 99.5% of the population as estimated from the available cohort. We could not build a population model for AMBP. (C) Extreme heterogeneity for AMBP individual dynamics with a very strong outlier. (D) Population dynamics, the gray area represents 95% of the population dynamics according to the model based on available data.

## CONCLUSION

In a previous publication, we introduced a mathematical model along with a data processing and filtering pipeline to analyze protein turnover data (Lehmann *et al*., 2019). This methodology, which we refer to as QNB in this paper, enabled us to obtain turnover parameters for individual proteins in individual samples. Here, we have proposed an extension to integrate the population level, *i*.*e*., the variability within a cohort of individuals for each protein. Population-level modeling relied on a hierarchical Bayesian approach combined with MCMC sampling to infer model parameters. This is an approach that is applied in population pharmacokinetics studies (Bauer *et al*., 2007; Duffull *et al*., 2005). Population protein turnover modeling enabled us to both describe inter-individual variability by means of the typical areas of the parameter space, and to derive accurate, individual-specific models able to infer her turnover parameters accurately. We also showed that the choice of a specific prior had no real impact on the posterior distribution due to sufficient experimental data.

Two MRM data sets were exploited. One human blood plasma data set featured limited inter-individual variability resulting in relatively easy modeling. Another human ventricular CSF data set featured high inter-individual variability, which challenged our methodology. We managed to obtain accurate models for 18/19 proteins available for every patient, the last case being pathological with one patient harboring massively different dynamics. On a larger cohort featuring high variability, one could imagine identifying sub-groups of patients beforehand, or including the notion of sub-group in the hierarchical Bayesian model, which is in principle well adapted for such tasks. This possibility will have to be explored in a future larger cohort.

This study has established a new type of mathematical model for the protein turnover community, which we believe should greatly facilitate the description and comparison of natural *versus* pathological protein turnover at the most relevant scale that is the population.

## SUPPLEMENTARY DATA

Tables S2-S5 are provided as supplementary data. Table S1 is in Supplementary Information (SI). All the individual protein data for all the patients are provided as supplementary data. Preprocessed input data, control plots, and the scripts implementing the Bayesian models are available from Zenodo (10.5281/zenodo.8425322). MRM data are available from https://panoramaweb.org/jGcF67.url. Blood plasma whole transcriptome data are available from https://panoramaweb.org/Xpd6J5.url. CSF whole transcriptome data were made public already (Lehmann *et al*., 2019).

## AUTHOR CONTRIBUTIONS

SL, CH, and JC conceived the study. JC and JPV analyzed the data. AG and LB provided patient material and clinical data. JV and CH analyzed the samples. JC, SL, CH, and JV wrote the manuscript. All authors have given approval to the final version of the manuscript.

## ACKNOWLEDGEMENTS

JC, SL, JPV, and CH were supported by the ANR-20-CE44-0007-01 grant. SL, AG, LB and CH were supported by the PHRC 2010 PROMARA. JC, SL, and CH were supported by the Fondation Alzheimer. JC thanks Olivier Gimenez for help starting with ODEs in BUGS.

