## Supplementary Information for "Enabling population protein dynamics through Bayesian modeling"

### SUPPLEMENTARY TABLES & FIGURES

**Table S1.** Demographic information (gray area irrelevant for plasma samples).

| Patient | Sex | Age | Diagnosis | Time after set up of ventricular drainage | CSF protein concentration (g/L) | Cell count per mm³ |
|---|---|---|---|---|---|---|
| Pat1a | F | 67 | Subarachnoid hemorrhage, Fisher scale 4, left carotid aneurysm | 19 days | 0.34 | 40 |
| Pat2a | F | 65 | Subarachnoid hemorrhage, Fisher scale 4, anterior communicating artery aneurysm | 15 days | 0.15 | 10 |
| Pat3a | F | 49 | Subarachnoid hemorrhage, Fisher scale 3, cerebral right posterior artery aneurysm | 23 days | NA | NA |
| Pat4a | F | 68 | Subarachnoid hemorrhage, Fisher scale 3, cerebral anterior artery aneurysm | 8 days | 0.50 | 150 |
| Pat7b | M | 57 | Fronto-temporal dementia | | | |
| Pat8b | F | 69 | Alzheimer's disease | | | |
| Pat9b | F | 49 | Mild Cognitive Impairment | | | |
| Pat10b | F | 80 | Alzheimer's disease | | | |
| Pat11b | M | 77 | Mild Cognitive Impairment | | | |
| Pat12b | F | 69 | Alzheimer's disease | | | |
| Pat13b | F | 61 | Alzheimer's disease | | | |

**Table S2.** Proteome-wide, unbiased analysis of Pat1 182 plasma proteins, turnover parameters. See Supplementary Data.

**Table S3.** Proteome-wide, unbiased analysis of Pat1 198 CSF proteins, turnover parameters. See Supplementary Data.

**Table S4.** Plasma proteins and peptides measured in MRM. See Supplementary Data.

**Table S5.** CSF proteins and peptides measured in MRM. See Supplementary Data.

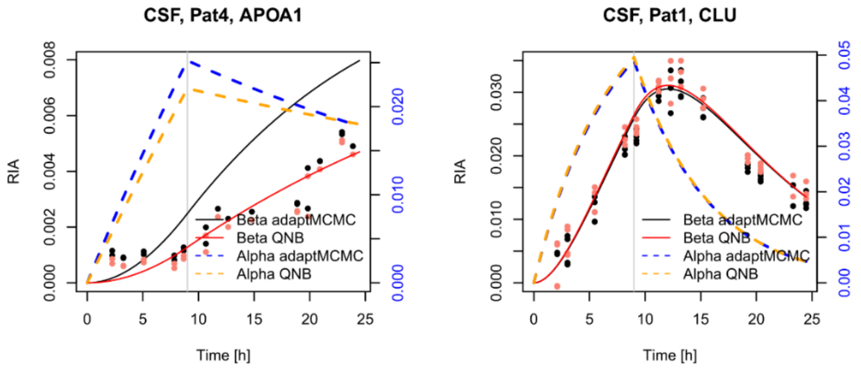


**Figure S1.** Failed and successful representative applications of adaptMCMC. (Left) An example where OpenBUGS managed to estimate model parameters leading to $\beta(t)$ going through the data points, while adaptMCMC inferred a similar shift but inadequate $\lambda$ and $k_c$. Data points are plotted in red and black according to the respective shifts of the two samplers. Here, 100 000 iterations with 50 000 burn-ins were used. To increase the number of iterations for adaptMCMC or to use its parallelized version with multiple chains did not improve, it always converged on this type of solution for data sets with this type of $\beta(t)$ geometry (data not shown). The same occurred with the library mcmc and its two samplers (data not shown). (Right) In the majority of cases, OpenBUGS and adaptMCMC or mcmc found very close and correct solutions as exemplified here with clusterin (CLU).

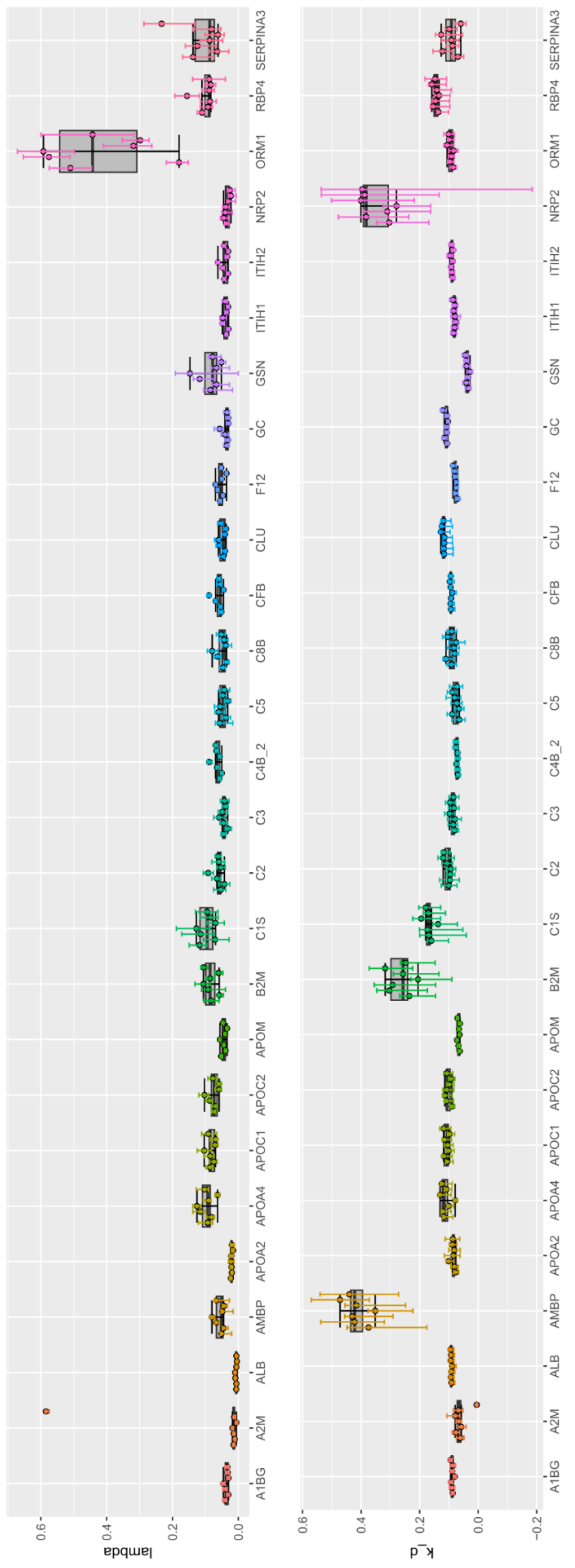


**Figure S2.** Blood plasma individual model parameter variability.

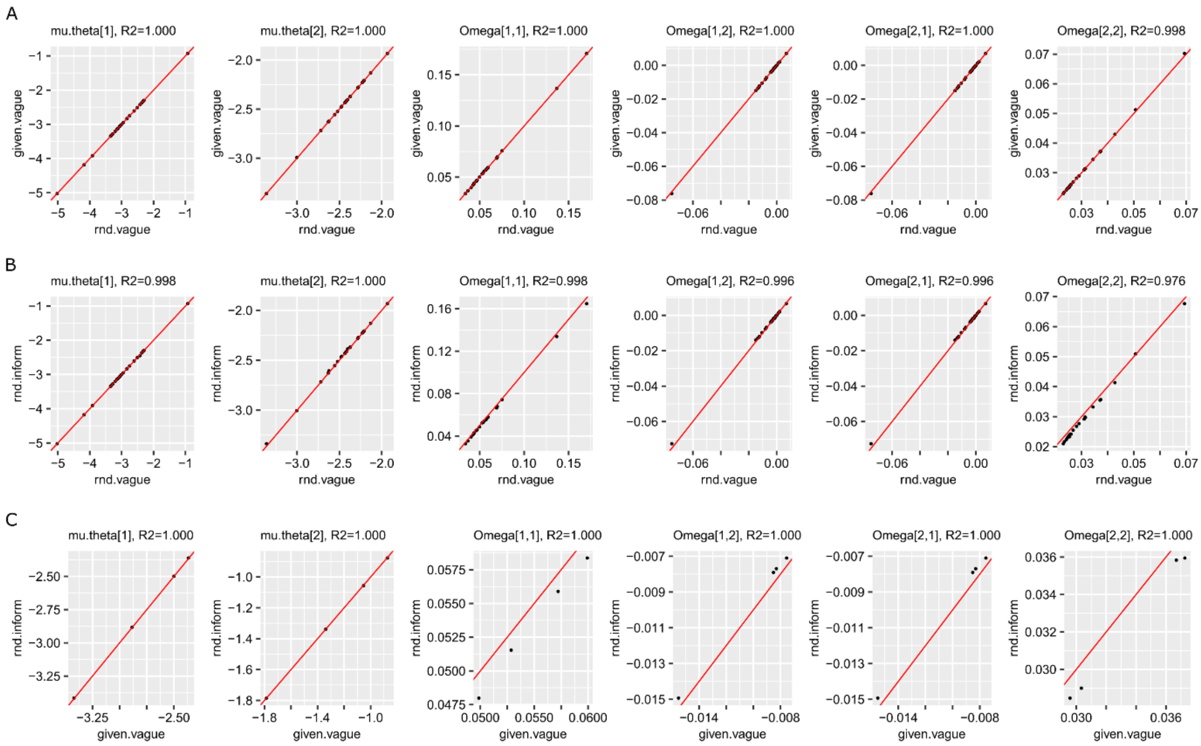


**Figure S3.** Comparison of the population model parameters obtained with the different approaches to start and run the MCMC sampling for the plasma proteins. (A) Random initial values *versus* those estimated by the QNB algorithm. Data from the 23 plasma proteins whose models were obtained with both approaches. (B) Random initial values *versus* more informative (hyper-)prior on $\mu_\theta$. Data from the 23 plasma proteins whose models were obtained with both approaches. (C) Comparison between the approaches with QNB initial values and a more informative prior for the four proteins (AMBP, B2M, C1S, and NRP2) whose model could not be obtained by the most basic and common approach (random initial values and vague prior).

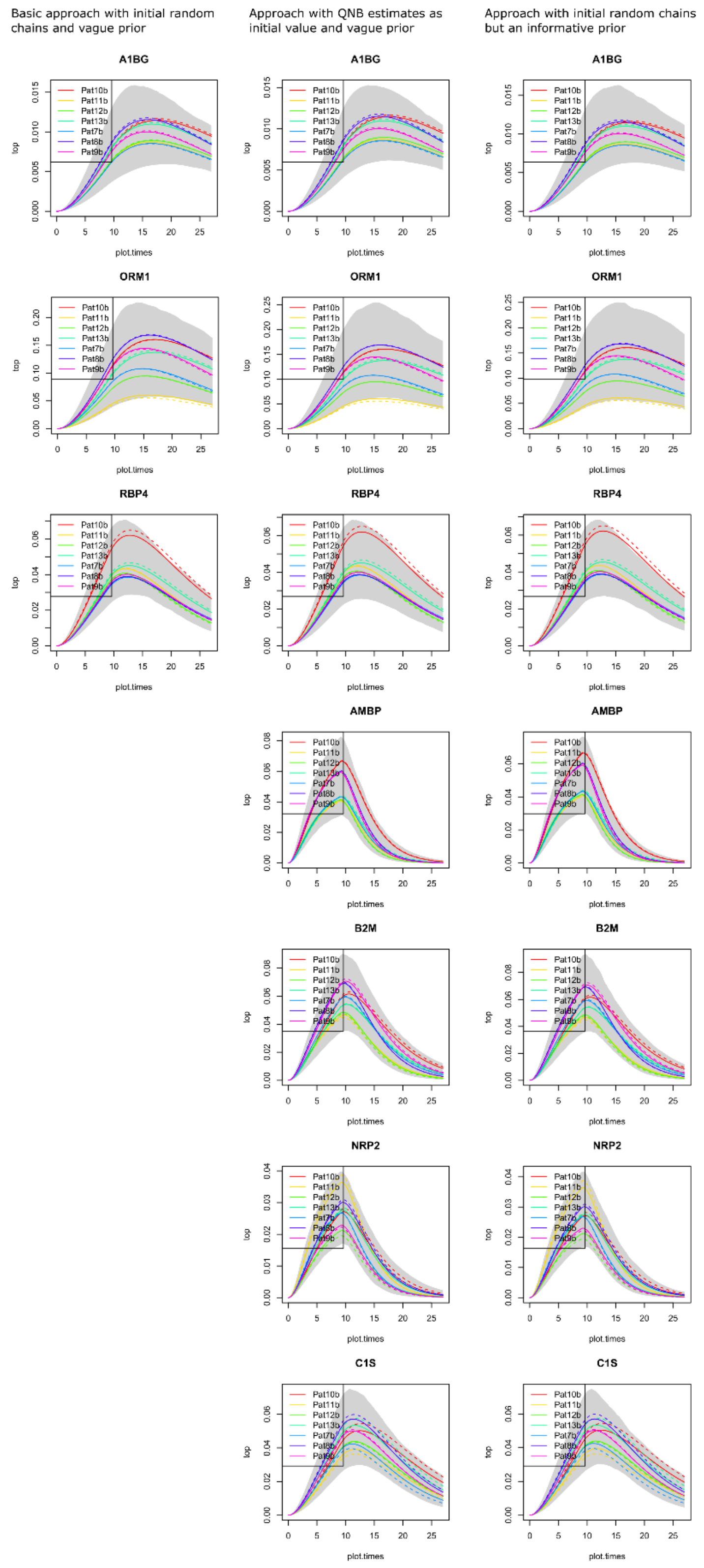


**Figure S4.** The population protein dynamics obtained with three approaches combining random or QNB given initial values for the Markov chains, or a vague *versus* a more informative (hyper-)prior on $\mu_\theta$. Three example plasma proteins whose population models were fit with all three approaches, followed by the four proteins whose models could not be obtained with the most basic and common approach.

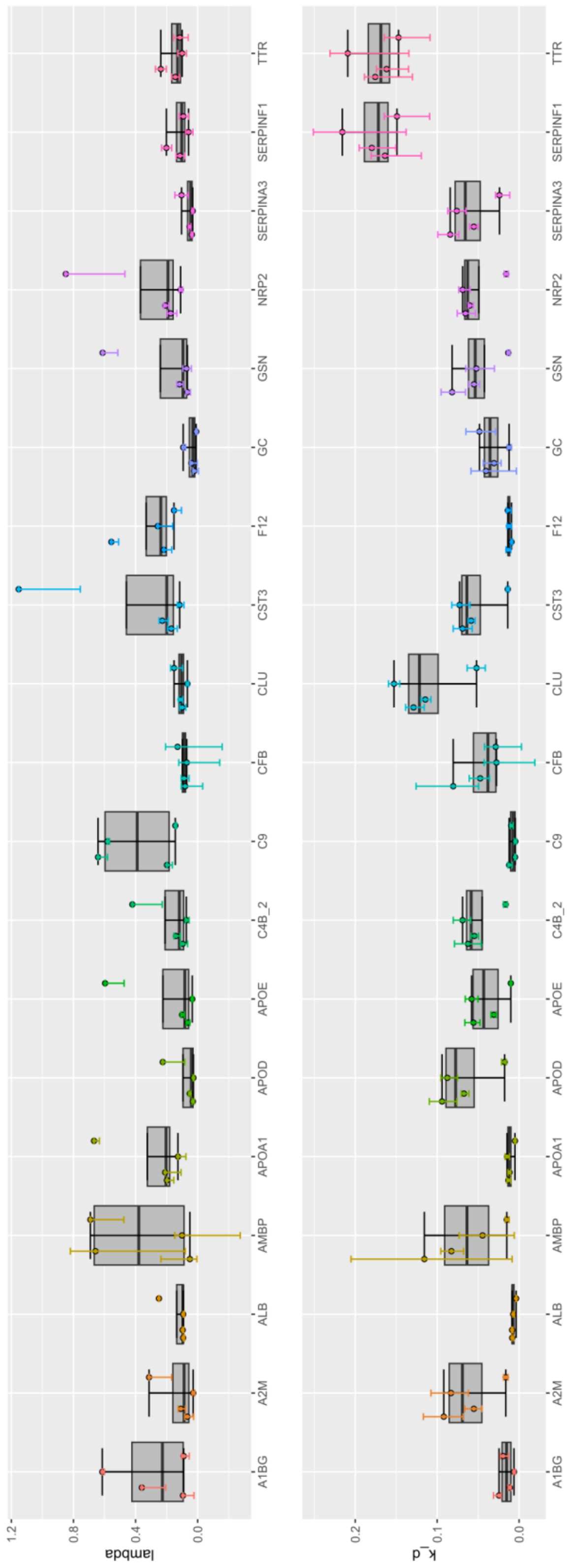


**Figure S5.** CSF individual model parameter variability.

### SUPPLEMENTARY METHODS

#### Individual protein BUGS model

```
# parameters: ngrid, ndim, init, times, origin, tol, llambda.mu,
# llambda.prec, lk_d.mu, lk_d.prec, shift.mu, shift.prec, weights,
# i_times

# ODE declaration
solution[1:ngrid,1:ndim] <- ode(init[1:ndim], times[1:ngrid],
                                D(C[1:ndim],t), origin,tol)

# ODE model priors
llambda ~ dnorm(llambda.mu, llambda.prec)
lambda <- exp(llambda)
lk_d ~ dnorm(lk_d.mu, lk_d.prec)
k_d <- exp(lk_d)
shift ~ dnorm(shift.mu, shift.prec)

# residuals and ODE definition
ilc <- step(9-t)
D(C[1], t) <- (lambda * ilc - C[1]) * k_d
D(C[2], t) <- k_d * (C[1] - C[2])
for (i in 1:n){
  j[i] <- i_times[i]
  mu[i] <- solution[j[i], 2] - shift
  w.tau[i] <- tau * weights[i]
  ratios[i] ~ dnorm(mu[i], w.tau[i])
}
# residuals prior
tau ~ dgamma(0.001,0.001)
```

**Likelihood function for adaptMCMC or mcmc R libraries**

```
# diff model ------------------------------

library(deSolve)

ilc <- function(t){
  if (t <= 9)
    1
  else
    0
}

model.2.1.j <- function(t,y,params){
  Alpha <- y[1]
  Beta <- y[2]
  dAlpha <- (params[1]*ilc(t)-Alpha)*params[2]
  dBeta <- params[2]*(Alpha-Beta)
  list(c(dAlpha,dBeta))
}
# likelihood

lupost <- function(par,ratios,pp){

  # parameters
  llambda <- par[1]
  lk_d <- par[2]
  shift <- par[3]
  sigma <- par[4]
  if (sigma<=0)
    return(list(log.density=-Inf))

  # log-prior
  lprior <- dnorm(llambda,pp$llambda.mu,llambda.sd,log=T) +
    dnorm(lk_d,pp$lk_d.mu,lk_d.sd,log=T) +
    dnorm(shift,shift.mu,shift.sd,log=T) +
    dgamma(1/sigma**2,shape=0.001,rate=0.001,log=T) # gamma prior for the
precision

  # log-likelihood
  times <- sort(unique(c(0,9,ratios$times)))
  parameters <- exp(par[1:2])
  ode.out <- radau(y=c(Alpha=0,Beta=0),times=times,func=model.2.1.j,
                  parms=parameters,events=list(time=9))
  LL <- 0
  for (i in 2:nrow(ode.out)){
    good <- ratios$times==ode.out[i,1]
    if (sum(good)>0)
      LL <- LL+sum(dnorm(ode.out[i,3]-ratios$ratios[good]-
shift,mean=0,sd=sigma,log=T))
  }

  list(log.density=LL+lprior)

} # lupost
```

**Population BUGS model for 7-patient plasma population**

```
# ODE system declaration
  solution[1:ngrid,1:ndim] <-
ode(init[1:ndim],times[1:ngrid],D(C[1:ndim],t),origin,tol)

# ODE parameters with their priors
for (i in 1:npat){
  lambda[i] <- exp(theta[i,1])
  k_d[i] <- exp(theta[i,2])
  theta[i,1:2] ~ dmnorm(mu.theta[1:2],inv.Omega[1:2,1:2])
}
# hyper-priors of the ODE parameters
mu.theta[1:2] ~ dmnorm(mu[1:2],T[1:2,1:2])
inv.Omega[1:2,1:2] ~ dwish(R[1:2,1:2],2)

# injection of heavy leucine
ilc <- step(9-t)

# ODE definitions
D(C[1], t) <- (lambda[1] * ilc - C[1]) * k_d[1]
D(C[2], t) <- k_d[1] * (C[1] - C[2])
D(C[3], t) <- (lambda[2] * ilc - C[3]) * k_d[2]
D(C[4], t) <- k_d[2] * (C[3] - C[4])
D(C[5], t) <- (lambda[3] * ilc - C[5]) * k_d[3]
D(C[6], t) <- k_d[3] * (C[5] - C[6])
D(C[7], t) <- (lambda[4] * ilc - C[7]) * k_d[4]
D(C[8], t) <- k_d[4] * (C[7] - C[8])
D(C[9], t) <- (lambda[5] * ilc - C[9]) * k_d[5]
D(C[10], t) <- k_d[5] * (C[9] - C[10])
D(C[11], t) <- (lambda[6] * ilc - C[11]) * k_d[6]
D(C[12], t) <- k_d[6] * (C[11] - C[12])
D(C[13], t) <- (lambda[7] * ilc - C[13]) * k_d[7]
D(C[14], t) <- k_d[7] * (C[13] - C[14])

# residuals
for (k in 1:npat){
  for (i in offset[k]:(offset[k+1]-1)){
    j[i] <- i_times[i]
    mu.resid[i] <- solution[j[i],2*k]-shift[k]
    w.tau[i] <- tau*weights[i]
    ratios[i] ~ dnorm(mu.resid[i],w.tau[i])
  }
  shift[k] ~ dnorm(shift.mu,shift.prec)
}
# residuals prior
tau ~ dgamma(0.001,0.001)
```
